# Transposon-plasmid nesting enables fast response to fluctuating environments

**DOI:** 10.1101/2025.06.04.657954

**Authors:** Yuanchi Ha, Rohan Maddamsetti, Xiaoli Chen, Emrah Şimşek, Dongheon Lee, Hyein Son, Charlotte Lee, Edo Kussell, Lingchong You

## Abstract

Mobile genetic elements (MGEs) play a critical role in shaping the response and evolution of microbial populations and communities. Despite distinct maintenance mechanisms, different types of MGEs can form nested structures. Using bioinformatics analysis of 14,338 plasmids in the NCBI RefSeq database, we found transposons to be widespread and significantly enriched on plasmids relative to chromosomes, highlighting the prevalence of transposon-plasmid nesting. We hypothesized that this nested structure provides unique adaptive advantages by combining transposition-driven genetic mobility with plasmid-mediated copy number amplification. Using engineered transposon systems, we demonstrated that nesting enables rapid and tunable responses of transposon-encoded genes in fluctuating environments. Specifically, transposition maintains a reservoir of the encoded genes, while plasmid copy number fluctuations further amplify the dynamic range of gene dosage, thus enhancing the response speed and stability of transposon-encoded traits. Our findings demonstrate an adaptive benefit of transposon-plasmid nesting and provide insights into their ecological persistence and evolutionary success.

## Introduction

Mobile genetic elements (MGEs) are fundamental drivers of microbial responses to environmental change by carrying genes that encode a wide variety of functions, including toxin degradation^1–5^, virulence^6–8^, and antibiotic resistance^9–14^.

Among the different types of MGEs, plasmids and transposons stand out due to their distinct but complementary functional roles^15,16^. Plasmids are extrachromosomal replicons that can replicate autonomously and are often maintained at variable copy numbers, allowing high gene expression, a property that is unique among MGEs^17,18^. They also serve as key vehicles for intercellular gene transfer, by conjugation or transformation^18,19^. Importantly, plasmid copy number (PCN) can vary dynamically, introducing dosage-dependent phenotypic heterogeneity within clonal populations^20–22^.

In contrast, transposons are DNA elements that mobilize genetic materials within their host via transposition^23,24^, often carrying cargo genes encoding critical traits, including antibiotic resistance and metabolic functions^25,26^. Moreover, a recent bioinformatic analysis of mobile genetic elements in prokaryotes found that antibiotic resistance genes are most significantly enriched in transposable elements^27^. Together, plasmids and transposons mediate many of the most consequential gene flow events observed in clinical and environmental settings^18,26–30^.

Despite their mechanistic differences, transposons and plasmids can often form nested structures^31–38^. Nesting refers to the physical insertion of one MGE into another, such as a transposon within a plasmid^32–35^, or an integron within a transposon^36–38^. This phenomenon has been reported across all types of MGEs, often in the context of resistance gene dissemination^27, 39–41^. These anecdotal and case-based studies suggest nesting may be both common and biologically important. For instance, the antibiotic resistance *bla_NDM_* gene, which is globally distributed across Gram-negative bacteria via more than 1,000 plasmids, first emerged on a transposon^33^. Moreover, layers of nesting could happen simultaneously. In one study, transposon Tn4401 carrying the antibiotic resistant *bla_KPC_* gene was found nested in a Tn2-like transposon, which was itself nested in different types of plasmids^35^.

While these examples suggest that nesting may enhance gene mobility, persistence, and phenotypic variability, the actual prevalence of nesting, and of transposon-plasmid nesting in particular, remains unclear. The recent bioinformatic analysis of mobile genetic elements in prokaryotes found that approximately 17% of transposase, the enzymes that facilitate transposon transpositions, are nested in other MGEs, including integrons, phages, phage-like element and conjugation element^27^. However, this analysis relies on recombinase-based classification, which lacks the resolution to distinguish conjugative elements and plasmids. As a result, the abundance of transposons on plasmids could not be determined. Thanks to the availability of a large amount of highly-quality genome sequences, we can now address this question more definitively^42^.

While the exact extent of transposon-plasmid nesting remains to be fully characterized, evidence suggests that it could offer two functional advantages. First, it enables intracellular and intercellular mobility of functional genes^43^. Under antibiotic selection, transposons can move from the chromosome onto plasmids, amplifying transposon-encoded gene copies within individual cells and across populations. If the plasmids are conjugative, these transposons can spread horizontally across different populations via plasmid transfer, thereby enabling broad dissemination of adaptive traits^34,35,43^. Second, transposon-plasmid nesting can amplify cell-to-cell variability in the expression of transposon-encoded genes^44,45^. If these genes encode antibiotic resistance, such variability contributes to heteroresistance – a phenotype characterized by subpopulations with variable resistance levels that complicate antibiotic treatment^44^.

In addition to facilitating gene flow and heterogeneity, transposon-plasmid nesting may also modulate temporal dynamics of the collective gene expression in a population. In particular, two studies cited above^43,44^ show that antibiotic selection drives an increase in the transposon-encoded antibiotic resistance genes, which is reversed upon antibiotic removal. This reversable modulation in response to environmental changes suggests a flexible tuning of the encoded genes. By itself, PCN can increase in the presence of positive selection of the plasmid and decrease when the selection is removed^20,46,47^. Moreover, such dynamic regulation is not limited to antibiotic resistance plasmids but is a more general property of plasmids. For instance, virulence plasmid copy number increases during the early expanding phase of the *Yersinia pseudotuberculosis* invasion and decreases in later phases^48,49^. Likewise, the transposon copy number (TCN) can be amplified by transposition in the presence of selection and reversed in the absence of section, including both antibiotic and environmental stress^50,51^. Therefore, the transposon-plasmid nesting could have a unique advantage as it can exploit an amplified dynamic range due to the two layers of regulation on gene dosage.

Building on these insights, we set out to test whether transposon-plasmid nesting provides advantages under fluctuating environments, such as those imposed by intermittent antibiotic exposures^43,44,46,47^. To this end, we developed a suite of engineered systems in *E. coli* containing different configurations of transposon-plasmid nesting, allowing independent tuning of transposition and plasmid copy number dynamics. These systems enable us to dissect the individual and combined contributions of transposition and PCN dynamics to the temporal dynamics of transposon-encoded genes. We found that transposon-plasmid nesting facilitates faster and more reliable adaptation under fluctuating conditions, with high-copy-number (HCN) plasmids showing especially rapid responses. Simulations suggest that this effect is driven by broader subpopulation variability in resistance gene dosage among cells harboring HCN plasmids. Taken together, these results support a quantitative framework for understanding the adaptive value of nesting and offer a new explanation for the evolutionary persistence of HCN plasmids, despite their metabolic burden.

## Results

### Transposons are enriched on plasmids

To quantify the occurrence of this nested structure, we conducted a bioinformatics analysis using on a previously curated dataset containing all bacterial and archaeal genomes in the NCBI RefSeq database with linked short-read sequencing data in the Sequencing Read Archive (SRA)^42^. This dataset consists of 14,338 plasmids from 4,317 bacterial and archaeal genomes. Since transposition requires enzymes called transposases, which bind to transposon ends to either copy or cut the transposon before pasting it into a new sequence, the presence of transposase sequences embedded within plasmid sequences serves as a reliable indicator of nested-MGE structures. We used this unique nested form of sequences to identify the occurrence of the transposon-plasmid nesting structures.

We first quantified the number of transposases per Mbp on both plasmids and chromosomes. Our findings show that transposases are commonly found on both chromosomes and plasmids but are five times more frequent per Mbp on plasmids than on chromosomes (**Figure 1A**). This normalized transposase copy comparison takes into account size difference between chromosomes and plasmids. Our results suggest that plasmids may serve as ‘jumping pads’ for transposons. We further showed that transposon-plasmid nesting is universal among plasmids of all copy numbers (**Figure 1B**). The inverse correlation between the PCN and proportion of plasmids that carry transposases can be explained by the inverse scaling between the PCN and plasmid length^42,52^. That is, low-PCN plasmids tend to be longer, and are thus more likely to contain more MGEs like transposons.

**Figure 1.**
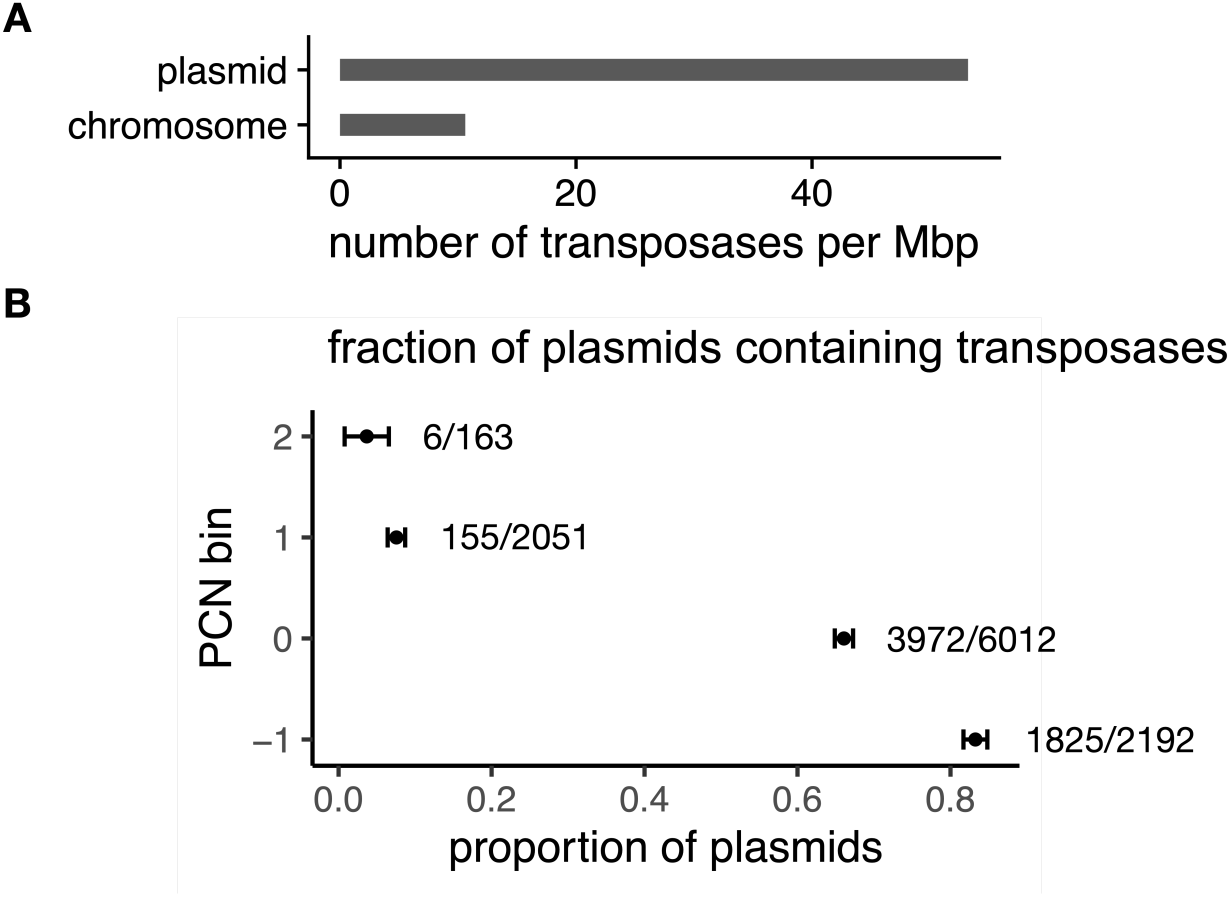
Transposon-plasmid nesting is highly prevalen. A. Bioinformatics analysis of the number of transposases per Mbp of 14,338 plasmids from 4,317 bacterial and archaeal genomes. Results show that transposons are five times more likely to be on plasmids than on chromosomes, when normalized by DNA length. B. Bioinformatics analysis of the existence of transposons on all plasmids, classified by the plasmid copy number (PCN). The y-axis is the log_10_(PCN), binned into 4 categories. The x-axis is the proportion of plasmids containing transposons in each category.

### Modeling predicted that plasmid-transposon nesting enabled fast transposon responses

We next sought to examine how transposon-plasmid nesting (**Figure 2A**) may contribute to the regulation of temporal dynamics of a transposon-encoded gene, by focusing TCN. We assume that each transposon contains a single copy of the target gene. Thus, TCN serves as a proxy for target gene dosage.

**Figure 2.**
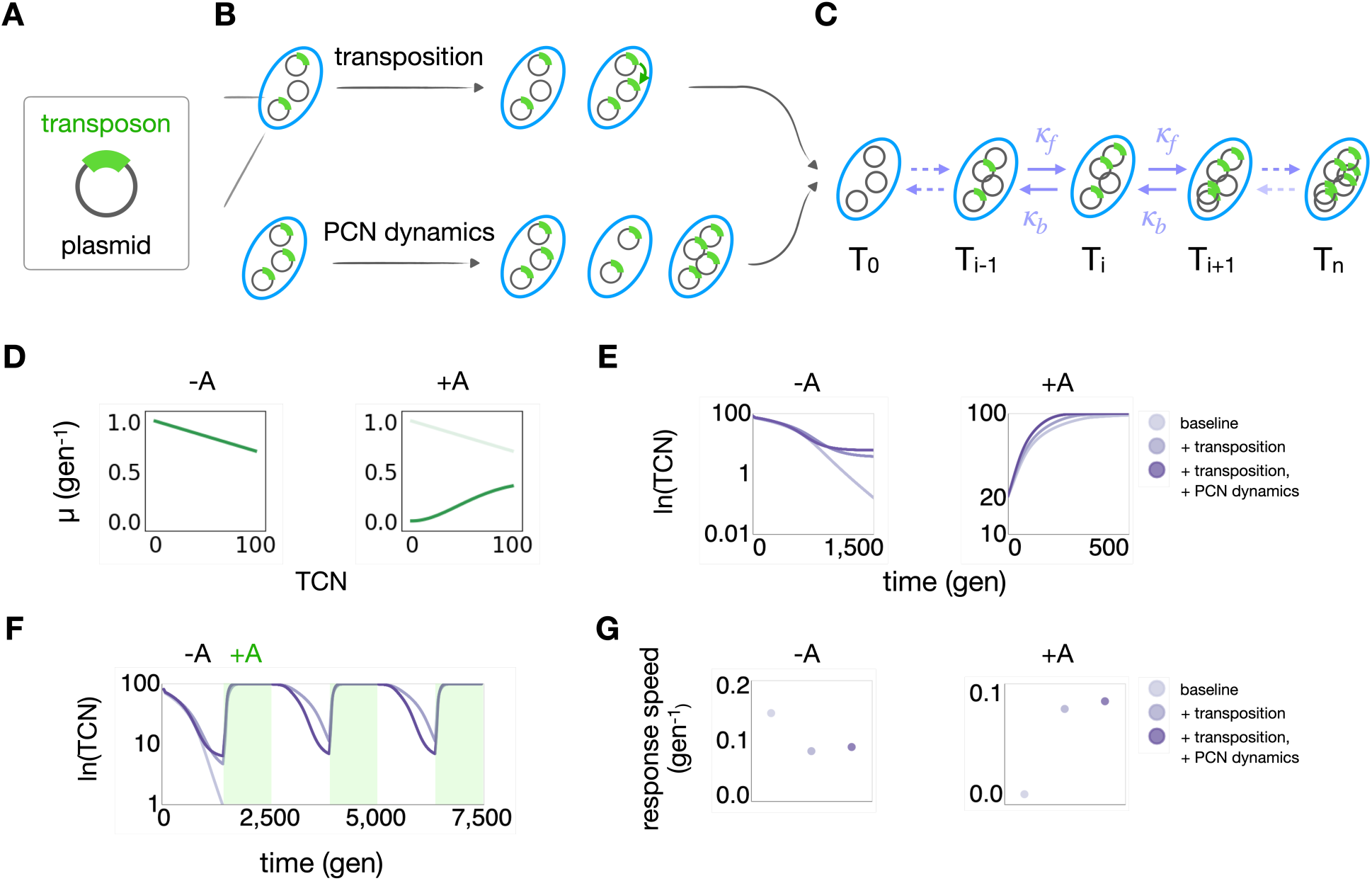
Modeling predicted that nesting enabled stable and fast transposon responses. A. Illustration of transposon-plasmid nesting. B. Illustration of how transposition increases the transposon copy number (TCN, top row) and how variations PCN drive those in the TCN (bottom row). C. Modeling framework. A subpopulation with TCN of *i* can transition to subpopulations *i* + 1 or *i* − 1, indicated by the arrows, and *T*_*i*_denotes the size of the *i*-th subpopulation. Transition rates are indicated by *k*_*f*_ and *k*_*b*_. D. Dependence of the growth rates ( *μ* ) of subpopulations carrying varying numbers of transposons. Without selection (left), we assume *μ* is a decreasing function of the transposon copy number. With selection, we assume *μ* increases with TCN following a Hill function. E. Simulated dynamics of TCN in fixed environments, under three different configurations: 1. Baseline: no transposition or PCN change (*k*_*f*_= 0.1, *k*_*b*_= 0.1); 2. With transposition but no PCN change (*k*_*f*_ = 0.15, *k*_*b*_= 0.1); 3. With both transposition and PCN change (*k*_*f*_ = 0.3, *k*_*b*_= 0.2). The left panel shows dynamics in the absence of selection; the right panel shows that in the presence of selection. Time is measured in units of generations. F. Simulated dynamics of TCN in fluctuating selection environments under the same three configurations. The third configuration (*k*_*f*_ = 0.3, *k*_*b*_ = 0.2) is the most responsive over time. G. Calculated response rates of all three populations. The baseline population has the highest loss rate and the lowest gain rate. Population with only transposition has the lowest loss rate. Population with transposition and plasmid copy number variation has higher loss rate than the second community and the highest gain rate overall.

We model a population where each cell can carry multiple plasmid copies, each of which may or may not contain a transposon. TCN per cell depends on both transposition (**Figure 2B**, top) and PCN dynamics (**Figure 2B**, bottom). Transposition increases the fraction of plasmids carrying the transposon, while PCN dynamics leads to cell-to-cell variation in TCN.

We use a coarse-grained framework to describe the temporal dynamics of subpopulations carrying varying TCNs (**Figure 2C**, **Methods**), as follows:

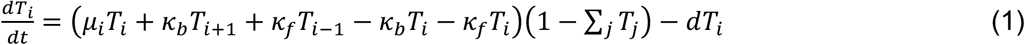

In this equation, *T*_*i*_ corresponds to the size of the subpopulation having *i* copies of the transposon, i.e. TCN = *i* ( 0 ≤ *i* ≤ *n*, where *n* is the maximum PCN) aggregated over subpopulations with different PCNs (Supplementary Figure 1). A system with *n* maximum plasmid copies consists of *n* + 1 subpopulations, each having a distinct TCN. Eq 1 models subpopulation growth, transition between subpopulations, and overall dilution.

We model the specific growth rate of *T*_*i*_ as 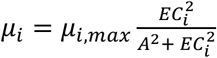, where *μ*_*i*,*max*_ is the maximum specific growth rate in the absence of antibiotic. We assume that *μ*_*i*,*max*_ decreases linearly with *i* to account for transposon burden (**Figure 2D**), while 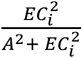 accounts for antibiotic tolerance by *T*_*i*_, where *A* is the antibiotic concentration and *EC*_*i*_is the antibiotic concentration needed to suppress *T*_*i*_growth by half. We assume *EC*_*i*_increases linearly with *i* (**Methods**). We assume logistic growth for all subpopulations population with a total carrying capacity of 1. Due to dilution (at rate *d*), the total population size will remain below the carrying capacity.

In our model, each transposition event increases TCN by one, corresponding to a transition *T*_*i*_ → *T*_*i*+1_ (**Figure 2B**, top). Given our experimental design, where there is a single copy of the transposase on the chromosome, we assume the rate of transposition is independent of the TCN. Upon cell division, the PCN is halved, and subsequently PCN increases over the duration of the cell cycle. Collectively, we refer to these processes as PCN dynamics (**Figure 2B**, bottom). Since we do not model individual cell division and plasmid replication events, the impact of PCN dynamics on TCN is incorporated in the model via the transition rates. Transitions *T*_*i*_ → *T*_*i*+1_ (with rate *k*_*f*_) and *T*_*i*_ → *T*_*i*–1_ (with rate *k*_*b*_) reflect gain or loss of a transposon, respectively, due to transposition or PCN dynamics. The transition *T*_*i*_ → *T*_*i*–1_ can happen due to PCN decrease after cell division, while the transition *T*_*i*_ → *T*_*i*+1_ can occur due to PCN increase after cell division or due to transposition. Even in the absence of transposition or PCN dynamics, there will be small, non-zero transition rates in both directions, resulting from random plasmid segregation during cell division, since cells cannot distinguish the plasmids carrying or not carrying the transposon in this process^53^. Thus, in the absence of transposition and PCN dynamics, we set *k*_*f*_ and *k*_*b*_to a low basal rate *k*_0_. Presence of transposition without PCN dynamics is represented by choosing *k*_*f*_ > *k*_0_and *k*_*b*_ = *k*_0_. Presence of both transposition and PCN dynamics is represented by further increasing *k*_*f*_, and choosing *k*_*b*_ > *k*_0_.

**Figure 2E** illustrates single simulated time courses of the same population with a maximum PCN of *n* = 100 and thus a maximum TCN of 100, with or without antibiotic selection (see **Supplementary Figures 2, 3** for simulations with a different value of *n*). We simulated three scenarios, corresponding to (1) no transposition or PCN dynamics ( *k*_*f*_ = *k*_*b*_ = 0.1 ), (2) transposition without PCN dynamics (*k*_*f*_ = 0.15, *k*_*b*_ = 0.1), and (3) both transposition and PCN dynamics (*k*_*f*_ = 0.3, *k*_*b*_ = 0.2). As **Figure 2E** shows, transposition promoted the maintenance of the transposon in the absence of selection and accelerated the TCN response in the presence of selection. That the exact value of *k*_*f*_ > 0.1 did not affect this conclusion (**Supplementary Figure 2**). As **Figure 2E** shows, transposition and PCN dynamics together accelerated TCN responses both in the presence and in absence of selection. The specific choices of parameter values *k*_*f*_and *k*_*b*_ did not affect the conclusion qualitatively (**Supplementary Figure 3**).

These results suggest that, over time, comparing to the baseline scenario where the cells have neither the transposition nor PCN dynamics mechanism, a population with transposition is more likely to maintain its transposons, and responds to changing environment faster with PCN dynamics. Continuously simulated time courses of these three populations indeed confirmed these two hypotheses (**Figure 2F**). For this simulation, we created a fluctuating environment where there are three cycles of antibiotic selection, each followed by a period of no antibiotic selection. The population with neither mechanism gradually lost all transposons over time. The same population with only the transposition mechanism stably maintained transposons. The population with both transposition and PCN variation further enabled faster transposon responses in both conditions (**Figure 2G**).

### Transposition promoted stable maintenance of the transposable element

To test model predictions, we adapted three transposon-plasmid pairs from our previous study^43^ (**Figure 3A**). The three plasmids are pBR322, CloDF13 and pUC. For brevity, we will call them pBR, CloDF and pUC in the rest of the paper. The three pairs differ by plasmid replication origin but have the same transposable element containing a tetracycline resistance gene (*tetA*) and a *gfp* reporter gene, as well as a kanamycin resistance gene (*kan*R) on the plasmid. The distinct antibiotic resistance markers on the plasmid and transposable element allow us to independently select for TCN (by using *tetA*) and PCN (by using *kanR*). We used GFP per unit biomass (GFP/OD600) as a surrogate measure of TCN, which we found to be consistent with, a more direct, qPCR-based quantification, except when TCN becomes too low (**Supplementary Figure 4**).

**Figure 3.**
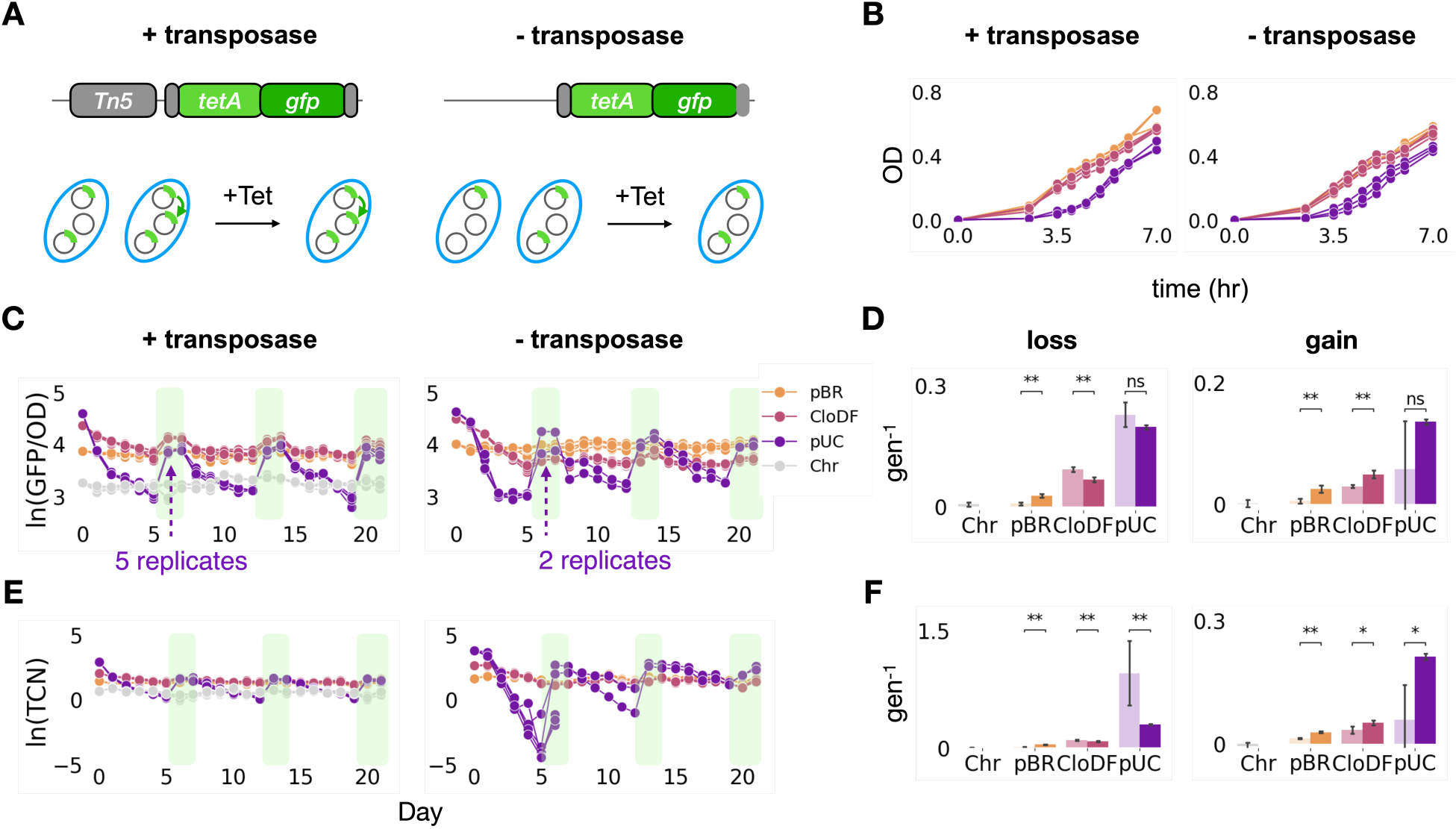
Experimental validation using synthetic systems of transposon-plasmid nesting. A. Illustration of the engineered system. B. The growth curves of strains containing different plasmids with transposase on the chromosome (left) or not (right). C. Temporal dynamics of OD-normalized GFP for strains containing different plasmids encoding the transposase (left) or not (right) in fluctuating environments. The identities of the plasmids are indicated by the three different colors and a different color, grey, indicates the strain that only carries the transposon on the chromosome. Note that without transposition, only two pUC replicates out of five can survive the tetracycline selection after the absence of tetracycline selection for the first five days. Legend is on the top right corner. D. Logarithm fold change of the OD-normalized GFP for both experiments. Left panel is for the loss rate and right panel is for the gain rate. Dark color indicates results of cells with transposase. Light color indicates results of cells without transposase. E. Temporal dynamics of TCN for both experiments. Data is presented in the same arrangement as in panel C. F. Logarithm fold change of the TCN for both experiments. Data is presented in the same arrangement as in panel D. Dark color indicates results of cells with transposase. Light color indicates results of cells without transposase. All statistical tests use the Mann-Whitney U test. p-value annotation legend: ns : 0.05< p <= 1.0; *: 0.01 < p <= 0.05; **: 0.001 < p <= 0.01

To examine the effect of transposition, we introduced these plasmids into two MG1655 hosts that differ in only one feature: one host has the transposase gene *Tn5* and the transposable element on the chromosome; the other does not. Therefore, one host contains functional transposons that are capable of transposition, and the other host contains unfunctional transposons. We exposed the different strains to periodic removal and addition of tetracycline, in three iterations for a total duration of 21 days, with five replicates per strain. Each iteration consisted of serial passaging of 30,000-fold dilution in fresh media, which corresponds to log_2_(30000) = ∼15 generations, without tetracycline for five days followed by two days with tetracycline (5 μg/mL), as the response to the tetracycline addition occurs more rapidly than that of following the removal of tetracycline. This tetracycline concentration was used as it allows the cells to grow for the full 15 generations per day, while also ensures the monotonic increasing max exponential growth rate from low-copy-number plasmid (pBR) to high-copy-number plasmid (pUC) under selection, aligning well with our simulation assumption (**Figures 2C, 3B**). We also added kanamycin (50 μg/mL) at each passaging to ensure PCN is maintained throughout the experiments, unless otherwise noted. Every ∼22 h, we measured GFP/OD600, stored samples, and performed a 30,000-fold dilution passage. Comparing the TCN changes of the two hosts (with or without a transposase) allowed us to determine the impact of transposon mobility.

The experimental results revealed significant differences in TCN loss and gain between cells with and without the transposase (**Figures 3C-F**). For cells carrying the highest copy number plasmid, pUC, the impact of transposon immobility was particularly striking. After the first five days without tetracycline selection, three out of five replicates failed to survive (OD < 0.05) subsequent tetracycline treatment (**Figures 3C, E**, right panels). The two surviving replicates retained extremely low TCN (ln(TCN)=-1 and −4, respectively), by the end of the first tetracycline removal period (**Figure 3E**). In contrast, cells with transposition all survived at the end of first tetracycline treatment and showed a shallower decrease in ln(GFP/OD) and in ln(TCN) over the five-day period without tetracycline selection. For cells carrying pBR and CloDF, even without transposition, all populations survived reintroduction of tetracycline selection and regained TCN.

We quantified the corresponding response rates by taking the averaged change of ln(GFP/OD) and ln(TCN) over the number of generations passed in between two treatments across all three fluctuating iterations (**Figures 3D, F, Methods**). For all three plasmids, the transposable elements were gained at a lower rate when there was no transposition, and for both CloDF and pUC plasmids, the transposable elements were lost at a faster rate when there was no transposition (**Figures 3D, F**), which confirmed with our simulation hypothesis. However, for pBR, the plasmid with the lowest copy number, the transposable elements were lost at a slower rate when there was no transposition. Upon further examination, we found that each pBR plasmid carried more than one transposon on average throughout the experiment. In contrast, each plasmid of the other two types, CloDF and pUC, carried less than one transposon (**Supplementary Figure 5**). Thus, the pBR population likely consisted of only plasmids that carried the transposons throughout the experiment. As such, it could not lose transposons during the fluctuating experiments. To further examine this notion, we did the same experiment with a MG1655 strain that only carried the transposase and the transposon on its chromosome and had no plasmids (**Figures 3C, E**). Since there was no plasmid for transposition or PCN dynamics in this setup, this population only consisted of one subpopulation. Indeed, it also had low rates of transposon gain or loss (**Figures 3D, F**).

This observed difference confirmed our model prediction that, in a population where not every plasmid already carries the transposon, transposition biases toward the subpopulations with higher TCNs by providing a continuous source for transposon generation. With transposition, each subpopulation has a higher probability of transitioning to a state with more TCNs, thus promoting subpopulations with higher TCN. Consequently, bacteria with transposition respond more rapidly under selective pressure and lose transposons more slowly in the absence of selection, compared with bacteria without transposition. Over extended periods, populations lacking transposition are more likely to lose all their transposons, rendering them unable to respond to similar stress in the future.

### Plasmid copy dynamics enabled faster transposon response

In the experiment above, the presence of the kanamycin selection throughout imposed a critical constraint on the dynamic range of PCN. In particular, it ensured that in each cell, the PCN was sufficiently high for a stable maintenance. We reasoned that removing kanamycin selection would remove this constraint and allow a larger dynamic range for the PCN dynamics.

As such, we repeated the experiment but without imposing the kanamycin selection (**Figure 4A**). Indeed, the PCN for each strain/plasmid combination exhibited a much larger dynamic range and faster responses (**Figures 4C, D, Methods**) in comparison with those in the presence of kanamycin selection. Concurrently, both GFP/OD600 and qPCR measurements showed faster TCN changes (**Figures 4E-H**) in comparison with those in the presence of kanamycin selection. The increases in the TCN response rates were more substantial for pBR (2.07-fold for gain rate, 2.35-fold for loss rate) and CloDF (2.14-fold for gain rate, 2.00-fold for loss rate) than for the pUC plasmid (1.04-fold for gain rate, 1.02-fold for loss rate). This could be due to the higher typical PCN of pUC. As such, the TCN on pUC has a wider range to adapt to fluctuating stress, i.e., from zero to the maximal PCN, via random segregation and transposition alone. In this case, the dynamic range of PCN was no longer the limiting factor. If so, we should see a faster TCN/PCN response.

**Figure 4.**
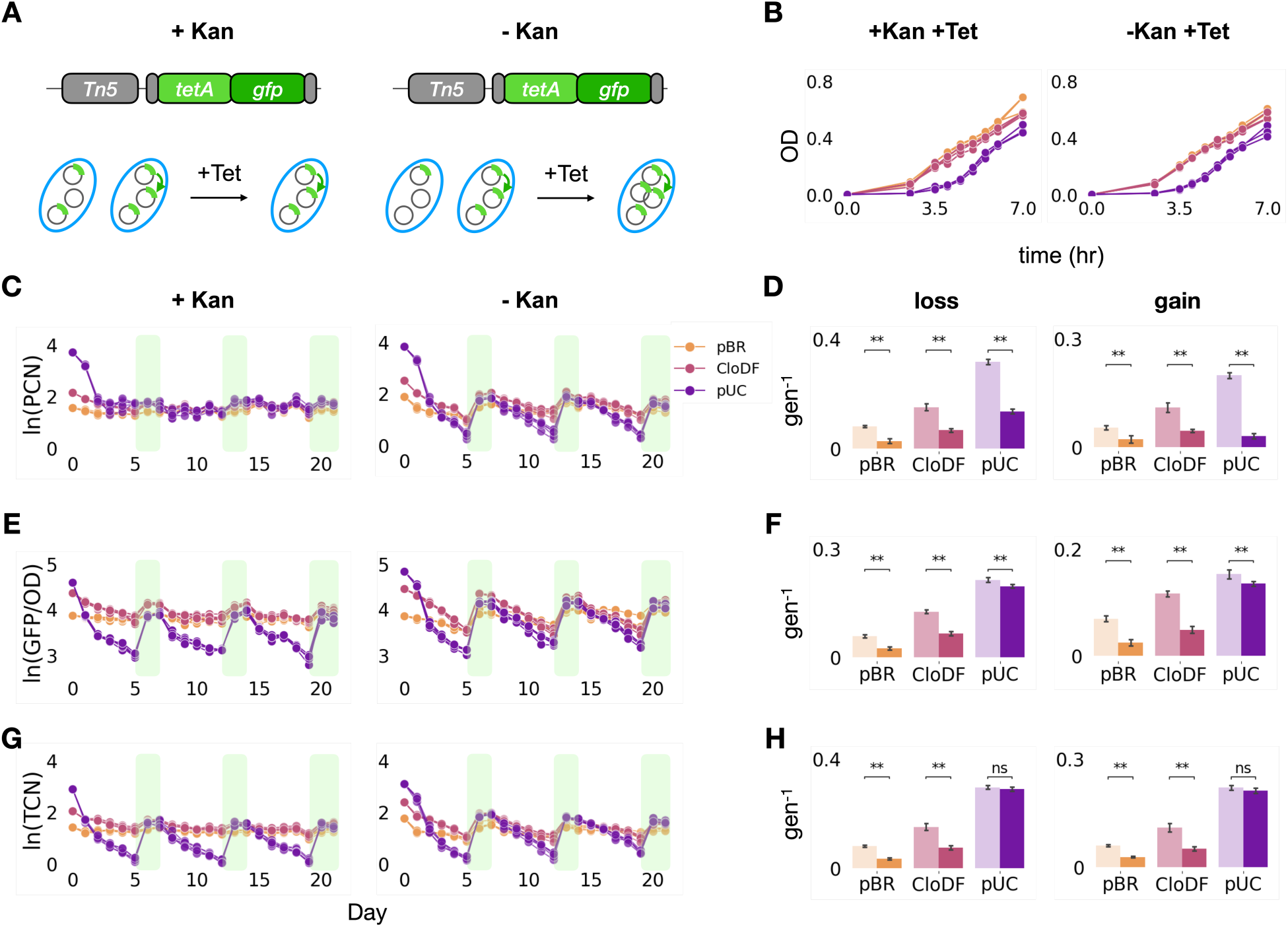
Plasmid responses amplified transposon responses. A. Experimental design. The kanamycin resistance gene allows for the control of the plasmid dynamic change in fluctuating tetracycline conditions. Kanamycin selection maintains the plasmids with reduced dynamic PCN variations (left); no kanamycin selection allows higher PCN variations (right). B. The growth curves of strains containing under both Kan and Tet selection (left) or under only Tet selection (right). C. Temporal dynamics of PCN for both experiments containing different plasmids under constant Kan selection (left) or not (right) in fluctuating environments. Left panel shows that constant application of kanamycin maintained the PCN in a narrow range. Right panel shows that PCN varied more drastically without kanamycin selection. The identities of the plasmids are indicated by the three different colors. Legend is on the top right corner. D. Logarithm fold change of the PCN for both experiments. Left panel is for the loss rate and right panel is for the gain rate. Dark color indicates results of cells under Kan selection. Light color indicates results of cells not under Kan selection. Cells with Kan selection gained and lost plasmids at slower rates. E. Temporal dynamics of OD-normalized GFP for both experiments containing different plasmids under constant Kan selection (left) or not (right) in fluctuating environments. Data is presented in the same arrangement as in panel C. F. Logarithm fold change of the OD-normalized GFP for both experiments. Data is presented in the same arrangement as in panel D. Dark color indicates results of cells under Kan selection. Light color indicates results of cells not under Kan selection. Cells with Kan selection gained and lost transposons at slower rates. G. Temporal dynamics of TCN for both experiments containing different plasmids under constant Kan selection (left) or not (right) in fluctuating environments. Data is presented in the same arrangement as in panel C. H. Logarithm fold change of the TCN for both experiments. Data is presented in the same arrangement as in panel D. Dark color indicates results of cells under Kan selection. Light color indicates results of cells not under Kan selection. Cells with Kan selection gained and lost transposons at slower rates. All statistical tests use the Mann-Whitney U test. p-value annotation legend: ns : 0.05< p <= 1.0; *: 0.01 < p <= 0.05; **: 0.001 < p <= 0.01

Our results confirmed this hypothesis (**Supplementary Figure 6**). The response rate of TCN/PCN was consistently higher in the presence of kanamycin treatment for plasmid retention for all three plasmids, but it was significantly more so for the pUC plasmid, with a five to ten-fold difference. Therefore, for transposon-plasmid with very high PCN, TCN/PCN and transient PCN dynamics can have similar effect on transposon response rates. Moreover, the TCN/PCN changed faster than the TCN without plasmid retention by kanamycin addition. This further confirms that, as the plasmids responded less to the tetracycline exposure, the TCN/PCN responded more to compensate for that constraint. This result shows that the plasmid amplification and TCN/PCN work together to respond rapidly under fluctuating environments.

We further hypothesized that the increased PCN response (in the absence of kanamycin selection) would promote TCN response rates, even in the absence of transposition and we did the same experiments on the strains that do not have the transposase on the chromosome. The result confirmed our hypothesis as well (**Supplementary Figure 7**). Moreover, bacteria harboring the transposon-plasmid pair in the highest copy number plasmid, pUC, did not recover under the tetracycline treatment following the first tetracycline free passage (**Supplementary Figures 7C, E**). This indicates that transposition in the nested plasmid-transposon pair ensures that the bacterial population retains its adaptive potential for the same kind of environmental challenge in the future.

Taken together, our results showed that while the transposition promotes the maintenance of the TCN, the PCN dynamic response amplifies the TCN response speed. The two mechanisms work together to ensure stable and fast transposon responses to fluctuating environments.

### HCN plasmids enabled faster population response in changing environments

Given the critical role of PCN dynamic range in promoting TCN response speed, we reasoned that a higher copy number plasmid on average will enable faster responses. We first tested this notion with simulations. To do so, we separately calculated the gain rate and the loss rate over PCN. When there is transposon selection, we ensured that the TCN at the starting point is close to the minimal TCN by assigning each population a starting composition of a normal distribution with a mean of 10 transposons and a standard deviation of 30% maximal transposons (see **Supplementary Figure 9A** for results of different means and standard deviations). On the other hand, when there is no transposon selection, since the maximal TCN for each population is different, we ensured that the TCN at the starting point is close to the maximal TCN by assigning each population a starting composition of a normal distribution with a mean of 90% maximal transposons and a standard deviation of 30% maximal transposons (see **Supplementary Figure 9B** for results of different means and standard deviations). To compare the response speeds of different PCN, we simulated populations of the maximal PCN of 10 to 100, with a discrete increase of 5 copies in between state. Indeed, the simulation results confirmed that higher PCN enables faster population responses in both environments (**Figure 5A**). Note the discontinuous jump in response speed as a function of PCN in the gain condition is due to the dilution rate being higher than the maximum growth rate of some low-PCN populations.

**Figure 5.**
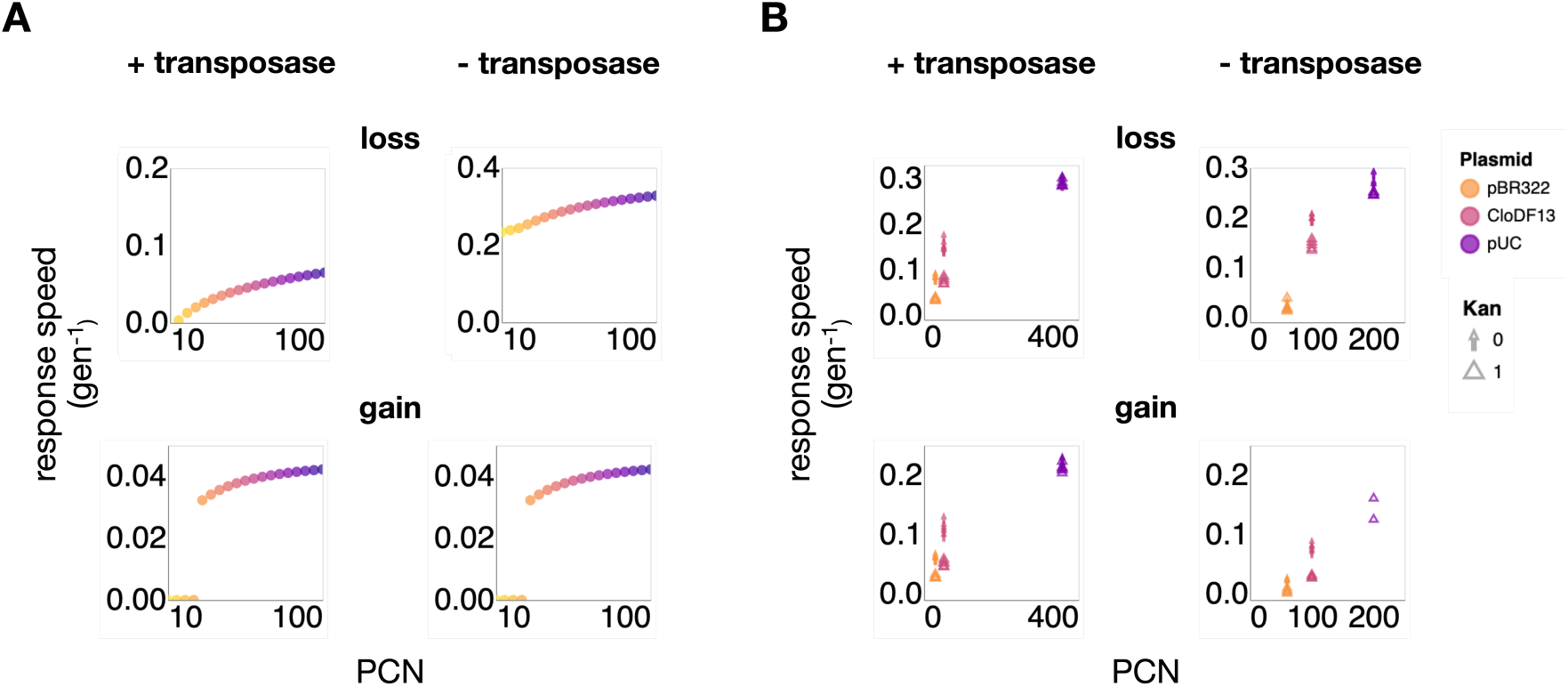
Plasmids with higher copy numbers enabled faster transposon response. A. Simulation analysis shows high-copy-number (HCN) plasmids enable faster population response. The simulation assumed higher PCN variations for HCN plasmids. The y-axis is the calculated response speed, and the x-axis is the maximal plasmid copy number. Color gradient also indicates the maximal PCN, with yellow indicating low copy number and purple indicating high copy number. B. Experimental data show HCN plasmids enable faster population response. The y-axis is the calculated response speed, and the x-axis is the maximal plasmid copy number. Top row is transposon loss rate; bottom row is transposon gain rate. The left side shows data of cells with transposon mobility; the right side shows data of cells with no transposon mobility. The arrows indicate the absence of Kan selection; triangles indicate the presence of Kan selection. Panel arrangement is the same as in panel A.

We think this result is due to the increasing competition between subpopulations as PCN, thus, the maximum TCN and subpopulation number of each population increase (**Figure 2D**). For example, the net growth rate difference between the subpopulations in a population containing only five subpopulations is much smaller compared to the net growth rate difference between subpopulations in a population with 100 subpopulations. Thus, stronger competition within the population of higher maximum PCN leads to faster dynamics in population response with more subpopulations. On the other hand, if the net growth rate difference ceases to increase as the PCN increases beyond a certain high value, the response rates cease to increase further after that point as well (**Supplementary Figure 10**).

Our experimental data indeed confirmed such positive relationship between the maximum PCN and the TCN response speed for all experimental conditions as well (**Figure 5B**). Plotting the transposon response rates with respect to the averaged PCN for a given plasmid revealed a strong positive correlation, both in the presence and absence of the tetracycline treatment. Averaged PCN here was calculated by averaging qPCR-based measurements of PCN over the entire experiment.

## Discussion

Our study reveals that transposon–plasmid nesting is both widespread and functionally significant. Transposons have long been known to move between bacterial genomes and MGEs and plasmids are recognized for their gene mobility and copy number variability. Here, we demonstrate that the nested structure creates a composite regulatory system with emergent properties. Specifically, this architecture enables stable maintenance and rapid modulation of gene copy number, allowing fast and robust responses by bacterial populations to fluctuating environments.

We showed that transposons are significantly enriched on plasmids compared to chromosomes. This enrichment spans all plasmid copy number classes and suggests that transposon-plasmid nesting is not restricted to specific MGE subtypes or ecological contexts. To dissect the functional consequences of this configuration, we used a coarse-grained model that tracks transitions between subpopulations with different TCNs. Simulations revealed that transposition primarily promotes gene persistence, while PCN variation accelerates gene dosage response. We validated these predictions using engineered bacteria, where PCN and transposition rate could be independently tuned. By minimizing confounding factors associated with natural systems, such synthetic systems are critical for uncovering the causal relationships between genetic architecture and population-level adaptation.

From an ecological perspective, the dual-layer regulation offered by nested MGEs may serve as a bet-hedging strategy: maintaining low-cost reservoirs of adaptive genes that can be rapidly amplified in a population under stress. The enhanced responsiveness conferred by high-copy-number (HCN) plasmids may also help explain their evolutionary persistence, despite the fitness costs they impose in the absence of selection. This mechanism contributes to explaining the longstanding “plasmid paradox” ^54^ --how costly plasmids persist over time --by showing that faster response speed under transient selection can offset the metabolic burden.

In fluctuating environments, such as those shaped by episodic antibiotic treatments, host immune pressure, or nutrient cycling, populations must rapidly adjust trait expression without losing the underlying genetic material. Transposon–plasmid nesting naturally supports this dynamic, encoding both memory and plasticity in the same construct. Importantly, this mechanism does not rely on prolonged positive selection, but instead exploits transient fluctuations, which are common in different types of microbial populations and communities^55,56^.

It has been suggested that one of the most far-reaching properties of plasmids is their multi-copy number per cell^17^. This property enables intracellular genetic variability. If the plasmids carry beneficial genes for the host, then such variability enables increased gene expression when the beneficial genes are selected and reduced gene expression when the selection is absent. Antibiotic resistance genes are among the most common plasmid-borne traits. It has been demonstrated that strong antibiotic selection increases PCN, therefore, enhances the antibiotic resistance, which is reversed after the selection is removed^45,46^. Likewise, transposon copies that carry beneficial genes can be amplified and reversed when the stress is introduced and removed^50,51^.

Our results also have implications for synthetic biology. Engineered gene circuits often suffer from a trade-off between stability (e.g., chromosomal integration) and flexibility (e.g., plasmid-based expression). A recent study has demonstrated that controlling plasmid loss rate, transfer rate and fitness effects, the fraction of plasmid-carrying cells and the plasmid-mediated gene expression can be amplified in the population level^21^. Transposon-plasmid nesting offers another layer for hybrid designs that encode both stability and fast, inducible responses. For instance, resistance genes or metabolic modules could be embedded within a transposon on transferable plasmids, enabling dynamic regulation of gene dosage in response to environmental inputs. Furthermore, our finding that transposon responses are faster in HCN plasmids suggests that PCN control could serve as a design lever for tuning system responsiveness without altering gene sequences or regulatory architecture. This is especially valuable in the design of community-based or fluctuating environment-responsive synthetic ecosystems^21,57,58^.

Finally, transposons have been found across the entire tree of life and have been observed nested not only in plasmids, but also in phages, integrons, and conjugative elements. We anticipate that the dual-layer control mechanism described here may generalize to other nested MGE architectures.

## Acknowledgement

This work was partially supported by the National Institutes of Health (L.Y., R01AI125604, R01GM098642, and R01EB031869; E.K., R01GM097356). The funders had no role in study design, data collection and analysis, decision to publish, or preparation of the manuscript.

## Author contributions

YH: conceptualization; investigation; methodology; resources; experimental data generation; data curation; formal analysis; visualization; validation; writing – original draft; project administration; writing – review and editing. RM: conceptualization; investigation; methodology; resources; bioinformatic analysis; experimental data generation; formal analysis; visualization; writing – review and editing. XC: qPCR experiments; investigation; writing – review and editing. ES: investigation; writing – review and editing. DL: validation. HS: validation. CL: supervision; writing – review and editing. EK: assistance with modeling analysis. LY: conceptualization; supervision; investigation; writing – original draft; project administration; writing – review and editing.

## Data availability

All data is available in the main text, supplementary information, or in GitHub at: https://github.com/youlab/transposon-plasmid-nesting

## Code availability

Code used for the computational pipeline described here and for figure generation is available in GitHub at: https://github.com/youlab/transposon-plasmid-nesting

## Methods

### Simulation framework

For a single population carrying one antibiotic resistant transposon and a plasmid of at most n copies, the population-level TCN change can be modeled as a set of n+1 ordinary differential equations (ODEs), each equation represents a subpopulation of a specific transposon number on plasmids:

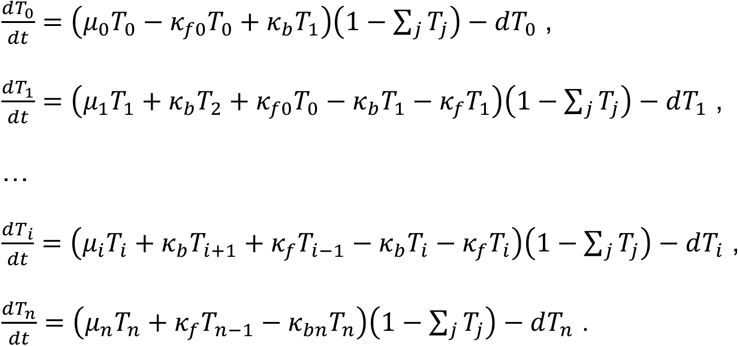

Here *T_i_* denotes the size of the subpopulation that carries *i* transposons on plasmids, 0 ≤ *i* ≤ n. *μ_i_* denotes the growth rate of that subpopulation. κ_f_ denotes the forward transition rate from one subpopulation to its neighboring population that has one more transposon; κ_b_ denotes the backward transition rate from one subpopulation to its neighboring population that has one less transposon. These two transition rates are the same between different subpopulations, except for the backward transition from *T_n_* to *T_n-1_* and the forward transition from *T_0_* to *T_1_*. Backward transition from *T_n_* to *T_n-1_* can happen when the transposase enzyme only cuts the transposable element but fail to insert it into a new location, which can occur occasionally. Therefore, we let *k*_*bn*_ = 0.1*k*_*b*_, assuming this event happens more rarely than the regular transposition event. Forward transition from *T_0_* to *T_1_* can only happen in our experimental setup where there is a transposon with the transposase on the chromosome to facilitate the transposition. Therefore, for the cells with transposition, *k*_*f*0_ = *k*_*f*_; for the cells without transposition, *k*_*f*0_ = 0.

To simulate the growth rate of each subpopulation under the two experimental conditions, i.e., one selects for and the other one selects against the transposons, the growth rate of each subpopulation is modeled in the following manner:

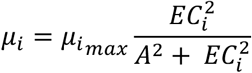

where

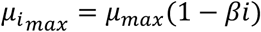

and

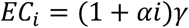

*μ_i_* is a function of both transposon burden and benefit. The left part *μ_imax_* models the transposon burden in a linear fashion. Specifically, subpopulations with increasing transposon copies have linearly decreasing growth rates by a constant factor β per additional transposon (Fig. 2C, left panel). However, when there is antibiotic selection, i.e., when A > 0, the subpopulation without resistant transposons, i.e., i = 0, has a growth rate of zero; and for subpopulations with increasing transposons, i.e., i > 0, the minimum inhibitory concentration (MIC) increases proportionally with i. This logic is modeled using a Hill equation 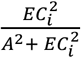 by the right part of the equation. Specifically, each additional transposon increases the half of the effective antibiotic concentration dosage γ by a constant factor α, thereby raising the antibiotic concentration, A, needed to reduce the growth rate by half. Consequently, under antibiotic selection, subpopulations with higher i exhibit increased growth rates in a Hill equation manner, capped by *μ_max_(1-βj)* (Fig. 2C, right panel).

### Strains, plasmids construction, and Transformation protocol

All experiments were conducted using *E. coli* strain MG1655. All plasmids were adopted from the previous publication^43^. We used electro competent^59^ cells for transforming all plasmids.

### Media and growth conditions

For all experiments, we cultured cells in LB media (LB broth Miller mix from Apex Bioresearch Products) at 37 °C at 225 rpm, supplemented with appropriate antibiotics. For the strain that only contain a transposon on the chromosome, without plasmid, the Tet concentration was 2 µg/mL. For the other strains, the Kan concentration was 50 µg/mL and Tet concentration was 5 µg/mL. For the long-term experiments, each day we first measured the OD and GFP, store the samples and did serial passaging. We first distributed (200 µl/well) of each sample into a black-walled 96-well plate (Corning, Cat. No. CLS3603). Cell density (optical density at 600 nm wavelength, or OD600) and GFP intensity (Ex: 488 nm, Em: 510 nm) were measured in a microplate reader (Tecan Infinite 200 PRO). Normalized GFP/OD was obtained by dividing GFP intensity by OD600 values. We then added 50 µl 50% mineral oil to each well and stored this 96-well plate in −80C for future qPCR quantification experiments. Overnight cultures were then diluted 3*10^−4^-fold into LB broth supplemented with appropriate antibiotics. To achieve this dilution fold, we did a two-step dilution by diluting 10-fold first in LB without antibiotics, then another 300-fold into LB with antibiotics.

### Growth rate calculation

For all strains, we grew them overnight in LB media at 37 °C at 225 rpm, supplemented with appropriate antibiotics, as mentioned in the above section. To measure the growth rate, we first diluted them 1000-fold the next day in fresh LB with the same antibiotic concentrations and let them grow four hours in the same condition until OD reaches 0.1. We then took 200 µl out every other hour to measure their OD600 values in a 96-well plate. We took the maximal OD difference per hour during the exponential phase as the growth rate for each strain.

### Primer amplification factor and primer efficiency calculation

qPCR primers and probes were designed to target (1) *cmR* or *dxs* gene on E. coli genome, for cells with transposase or without transposase (2) *kanR* gene on plasmid and (3) *tetA* gene on transposon. The *cmR* and *dxs* genes are present approximately one copy per genome, enabling us to calculate the average per-cell transposon and plasmid copy numbers relative to the genome copy number. An overnight culture of *E. coli* MG1655 carrying the plasmid was used to calculate the amplification factor and primer efficiency for each primer pair. The overnight culture was diluted by mixing 50 μl of culture with 150 μl of molecular-grade water. For cell lysis, 50 μl of the diluted culture was boiled at 98°C for 20 min. Then cell lysate was serially diluted (1/10-fold) and 4 µl of each diluted sample was used in qPCR reactions, with 6 technical replicates performed for each dilution rate. qPCR reactions were thermal cycled at 95 °C for 20 seconds, followed by 40 cycles of 95 °C for 1 second and 60 °C for 20 seconds, using TaqMan™ Fast Advanced Master Mix (Applied Biosystems, Cat. No. 4444965) in QuantStudio™ 5 Real-Time PCR system (Applied Biosystems).

Ct values were plotted against log-transformed dilution rates (**Supplementary Figure 8**), followed by linear regression to obtain the slope of the standard curve. The amplification factor was calculated using the formula: 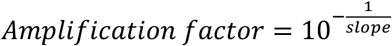. The primer efficiency was then derived as: *Primer efficiency* = (*amplification factor* − 1) × 100. Primer efficiencies of 100.91 % (amplification factor = 2.01), 103.47 % (amplification factor = 2.03) and 98.97 % (amplification factor = 1.99) were determined for *cmR, kanR* and *tetA* binding primers, respectively, for cells with transposase. Primer efficiencies of 97.46 % (amplification factor = 1.97), 100.57 % (amplification factor = 2.01) and 95.14 % (amplification factor = 1.95) were determined for *dxs, kanR* and *tetA* binding primers, respectively, for cells without transposase.

### qPCR quantification

Each day, we used the same samples used for OD and GFP measurements to quantify average plasmid copy number and transposon copy number to genome. The overnight culture was diluted by mixing 50 μl of culture with 150 μl of molecular-grade water. For cell lysis, 50 μl of the diluted culture was boiled at 98°C for 20 min. Then cell lysate was serially diluted (1/10-fold) and 4 µl of each diluted sample was used in qPCR reactions, with 3 technical replicates performed for each dilution rate. qPCR reactions were thermal cycled at 95 °C for 20 seconds, followed by 40 cycles of 95 °C for 1 second and 60 °C for 20 seconds, using TaqMan™ Fast Advanced Master Mix (Applied Biosystems, Cat. No. 4444965) in QuantStudio™ 5 Real-Time PCR system (Applied Biosystems).

Average plasmid copy number relative to genome was calculated by: 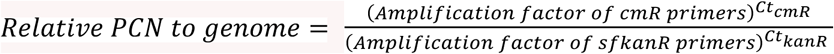 for cells with transposases, and 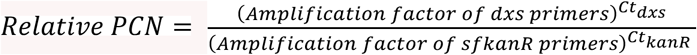 for cells without transposases, where Ct_dxs_, Ct_cmR_ and Ct_kanR_ correspond to cycle thresholds for *dxs*, *cmR* and *kanR* targets, respectively.

Average transposon copy number relative to genome was calculated by: 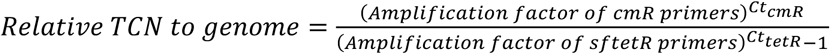 for cells with transposases, and 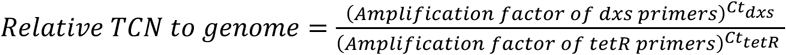 for cells without transposases, where Ct_dxs_, Ct_cmR_ and Ct_tetR_ correspond to cycle thresholds for *dxs*, *cmR* and *tetR* targets, respectively. Note we subtract the one transposon copy on the chromosome here for the cell with transposases for the relative TCN to genome calculation, since the one transposon copy we put on chromosome will not get lost during transposon responses.

### Response speed calculation

For the plasmid or transposon response rates, we averaged the response rates from all three fluctuating treatments. We calculated the response rates using a growth rate formula, which is the log change of PCN or TCN per generation. Using the transposon response rates after the first seven day treatment as an example, since the bacteria was subjected to no transposon selection of five days followed by selection of two days and each day, we diluted the cell 30,000-fold, the gain rate is calculated as 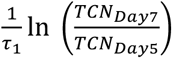, where *τ*_1_ = 2 log_2_ (30,000) generations, and the loss rate is calculated as 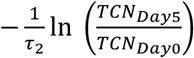, where *τ*_2_ = 5 log_2_ (30,000) generations. We used similar formulas for simulation results 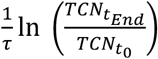, where 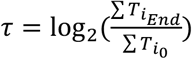 generations. Here 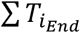 is the final population density by summing the densities of all subpopulations at the final simulation time point, and 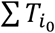 is the initial population density by summing the densities of all subpopulations at the initial simulation time point.

