## Supplementary Table and Figures for "Transposon-plasmid nesting enables fast response to fluctuating environments"

**Table S1: Model parameters**

The parameters used in the modeling were non-dimensionalized (ND) with respect to the carrying capacity and maximum growth rate. We used following parameters.

| Parameter name | Description | ND Value(s) (value ranges for simulation) |
|---|---|---|
| $\mu_{max}$ | Maximal growth rate of transposon-free subpopulations | 1 |
| $\beta$ | Burden from carrying each transposon | 0.003 |
| $\alpha$ | Benefit from carrying each transposon | 30 |
| $\gamma$ | Half of the minimal inhibition concentration of the antibiotics | 0.01 |
| A | Antibiotic concentration | 2 |
| d | Dilution rate | 0.05-0.3 |
| $\kappa_f$ | Forward transition rate | 0.1-0.4 |
| $\kappa_b$ | Backward transition rate | 0.1-0.2 |

**Supplementary Figure 1. Simplified model assumptions**

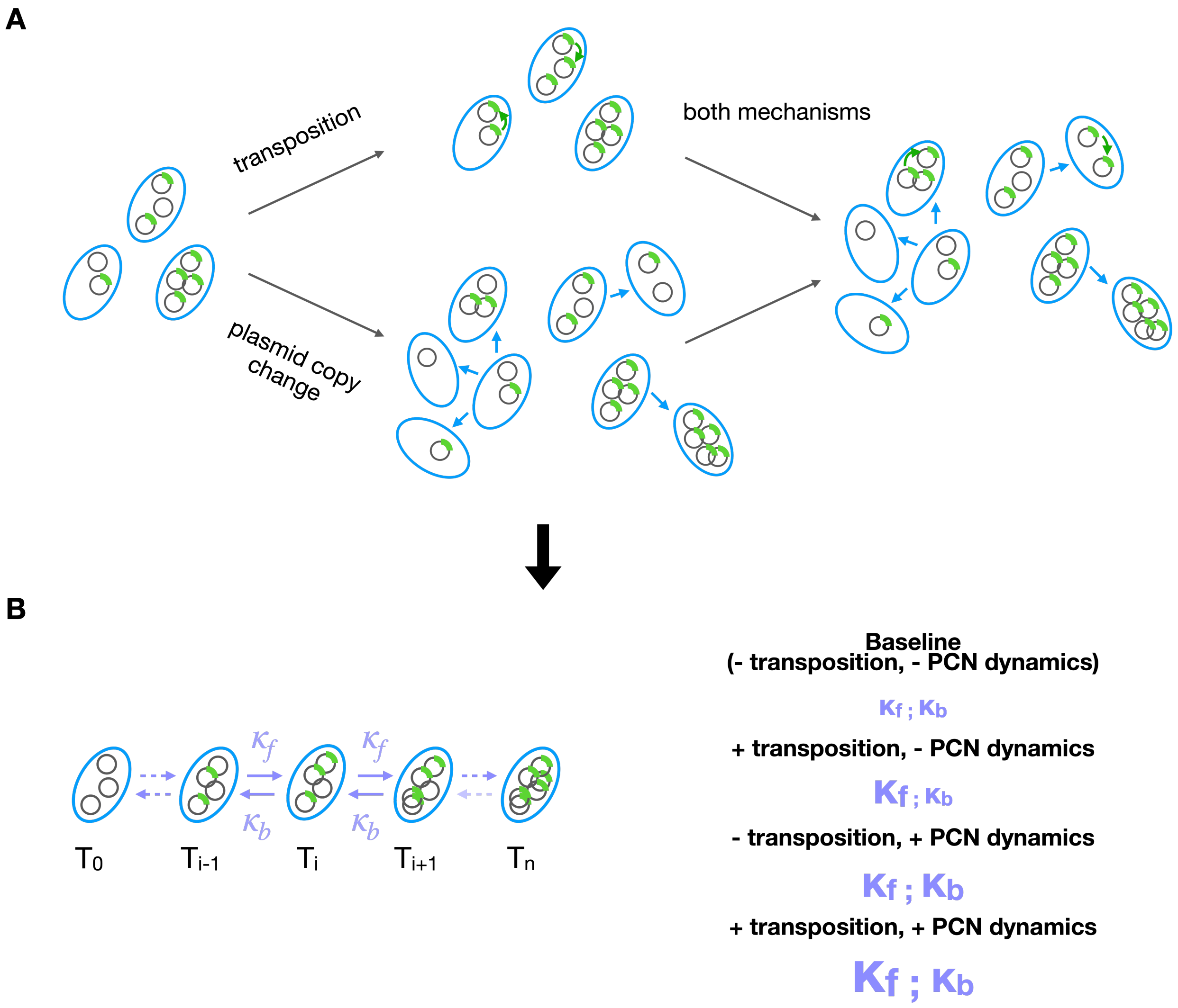


A. Detailed illustration of how transposon-plasmid nesting utilizes both transposon transposition and plasmid copy number dynamics mechanisms to achieve higher overall transposon copy number and larger transposon copy variation in a population started with different subpopulations. The left panel is the base case where there is no transposon transposition or plasmid copy number dynamics. The bacteria population still maintains subpopulations of different transposon copies due to plasmid random segregation. The middle panel shows the two mechanisms individually. The top middle panel shows that the transposon transposition mechanism increases the overall transposon copies in each subpopulation. The bottom middle panel shows that the plasmid copy number dynamics increases the overall transposon copy variation. The blue arrows indicate plasmid copy number dynamics. The right panel is the case where both mechanisms are in play.

B. Simplified model where only transposon copy numbers are explicitly modeled. The left side is an illustration of the model, where one population of at most n plasmids have n+1 subpopulations, each carrying one specific number of transposons. Each subpopulation could transition to its one or two neighboring subpopulations, indicated by the arrows in between. The forward transition rate is indicated by $\kappa_f$ and the backward transition rate is indicated by $\kappa_b$. The right side is an illustration of the representations of the four cases using our model, where each of the transposon transposition or plasmid copy plasmid copy number dynamics mechanism may or may not exist. The font size of the transition rates indicates their relative values in all four scenarios.

**Supplementary Figure 2. Simulation results assuming different effects of transposition on $\kappa_f$ show the same trend**

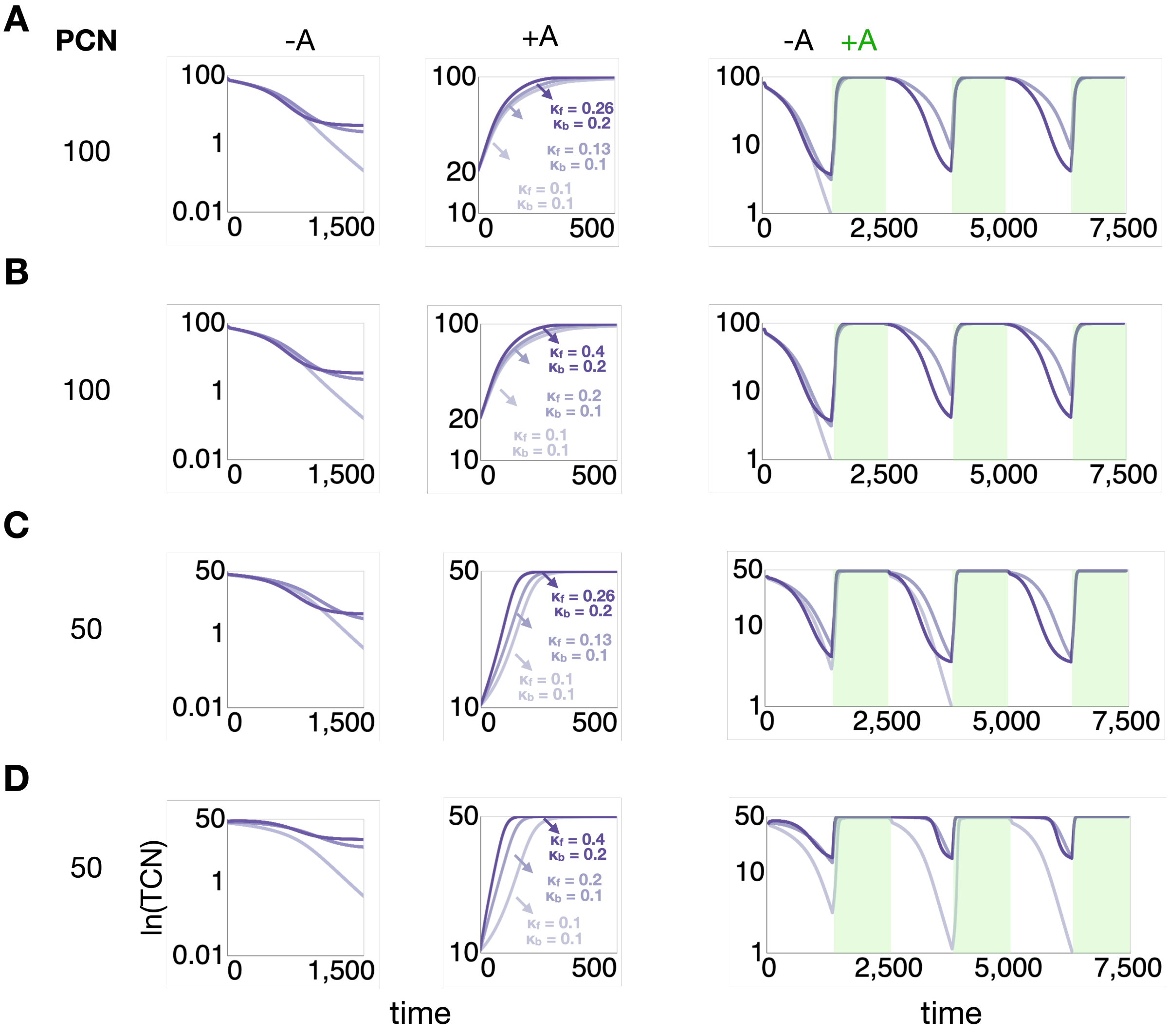


Each row contains simulation results of both single time courses and fluctuating environments.

A-B. Simulated population of 100 maximal PCN, thus 50 maximal TCN on plasmids.

C-D. Simulated population of 50 maximal PCN, thus 50 maximal TCN on plasmids.

A. Transposition increases $\kappa_f$ from 0.1 to 0.13, without the PCN dynamics effect.

B. Transposition increases $\kappa_f$ from 0.1 to 0.2, without the PCN dynamics effect.

C. Transposition increases $\kappa_f$ from 0.1 to 0.13, without the PCN dynamics effect.

D. Transposition increases $\kappa_f$ from 0.1 to 0.2, without the PCN dynamics effect.

In all cases, transposition helps to maintain transposons when there is no selection and elevates transposon copies when there is selection; plasmid copy response increases transposon response speeds both under no selection and under selection.

**Supplementary Figure 3. Simulation results assuming different effects of PCN change on $\kappa_f$ and $\kappa_b$ show the same trend**

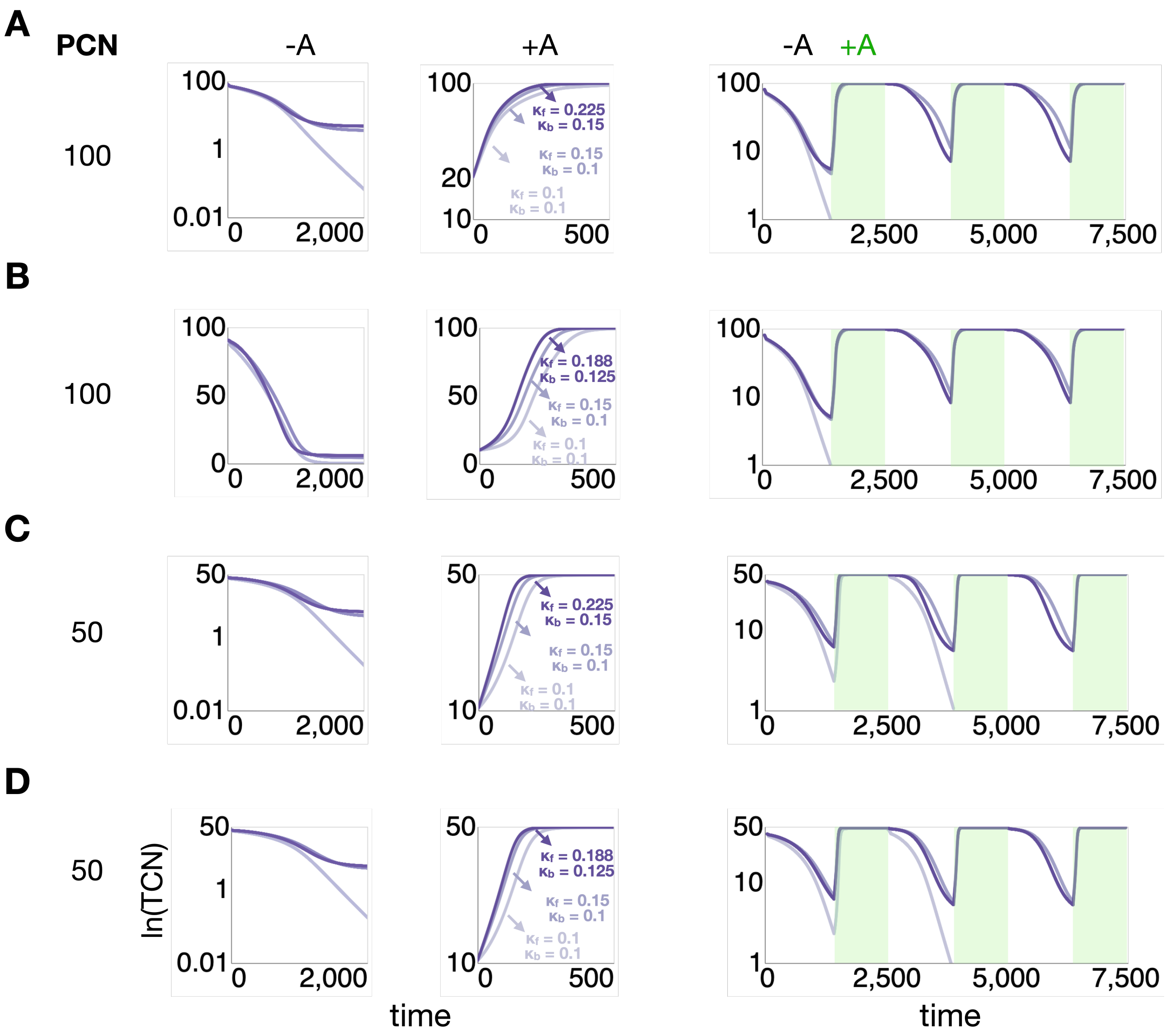


Each row contains simulation results of both single time courses and fluctuating environments.

A-B. Simulated population of 100 maximal PCN, thus 50 maximal TCN on plasmids.

C-D. Simulated population of 50 maximal PCN, thus 50 maximal TCN on plasmids.

E. PCN dynamics increases $\kappa_f$ and $\kappa_b$ by 1.5-fold.

F. PCN dynamics increases $\kappa_f$ and $\kappa_b$ by 1.25-fold.

G. PCN dynamics increases $\kappa_f$ and $\kappa_b$ by 1.5-fold.

H. PCN dynamics increases $\kappa_f$ and $\kappa_b$ by 1.25-fold.

In all cases, transposition helps to maintain transposons when there is no selection and elevates transposon copies when there is selection; plasmid copy response increases transposon response speeds both under no selection and under selection.

**Supplementary Figure 4. GFP/OD well represented transposon copy number from qPCR quantification**

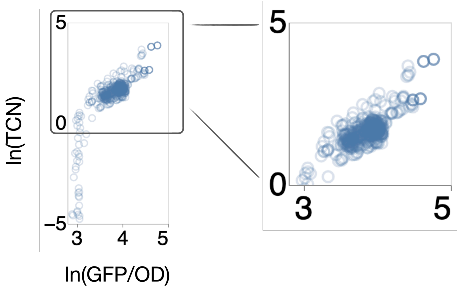


Left panel is the scatter plot of ln(TCN) from qPCR quantification versus ln(GFP/OD) from all experimental results. The right panel is a zoomed in section of ln(TCN) >= 0. ln(TCN) < 0 indicates there is less than one transposon per cell, thus, extremely low GFP/OD that could not be detected, which explains the vertical line from -5 <= ln(TCN) < 0.

**Supplementary Figure 5. TCN/PCN data for cells without transposase when PCN dynamics was limited in Fig. 3**

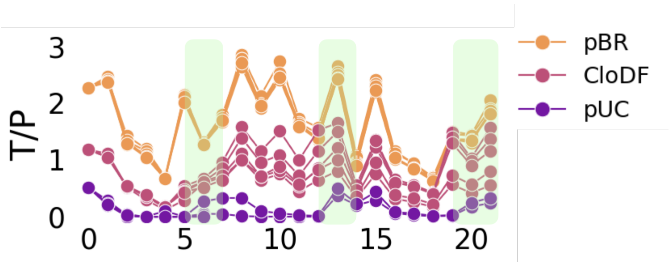


Each pBR plasmid carries more than one transposon on average throughout the experiment. Each CloDF or pUC plasmid carries less than one transposon on average throughout. This explains why transposon-pBR plasmid pair cannot lose the transposable element as efficiently when there is no transposase.

**Supplementary Figure 6. Dynamic responses were mediated by changes in TCN/PCN when PCN dynamics was limited**

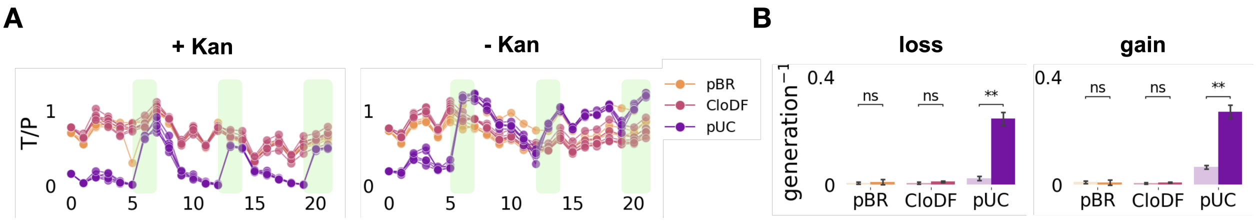


When there is transposition, high copy number plasmid, pUC shows large TCN/PCN change when there is limited PCN change.

A. qPCR results of the TCN per PCN (TCN/PCN) for both conditions, under constant Kan selection and under no Kan selection, of all three plasmids.

B. Logarithm fold change of (TCN/PCN) for both conditions. Dark color indicates results with Kan selection. Light color indicates results without Kan selection. Dark bars are higher than light bars and significantly so for pUC plasmid, indicating that when PCN dynamics is limited by Kan selection, TCN/PCN responds more.

All statistical tests use the Mann-Whitney U test. p-value annotation legend:

ns : $0.05 < p <= 1.0$; *: $0.01 < p <= 0.05$; **: $0.001 < p <= 0.01$

**Supplementary Figure 7. PCN dynamics accelerated transposon response without transposase**

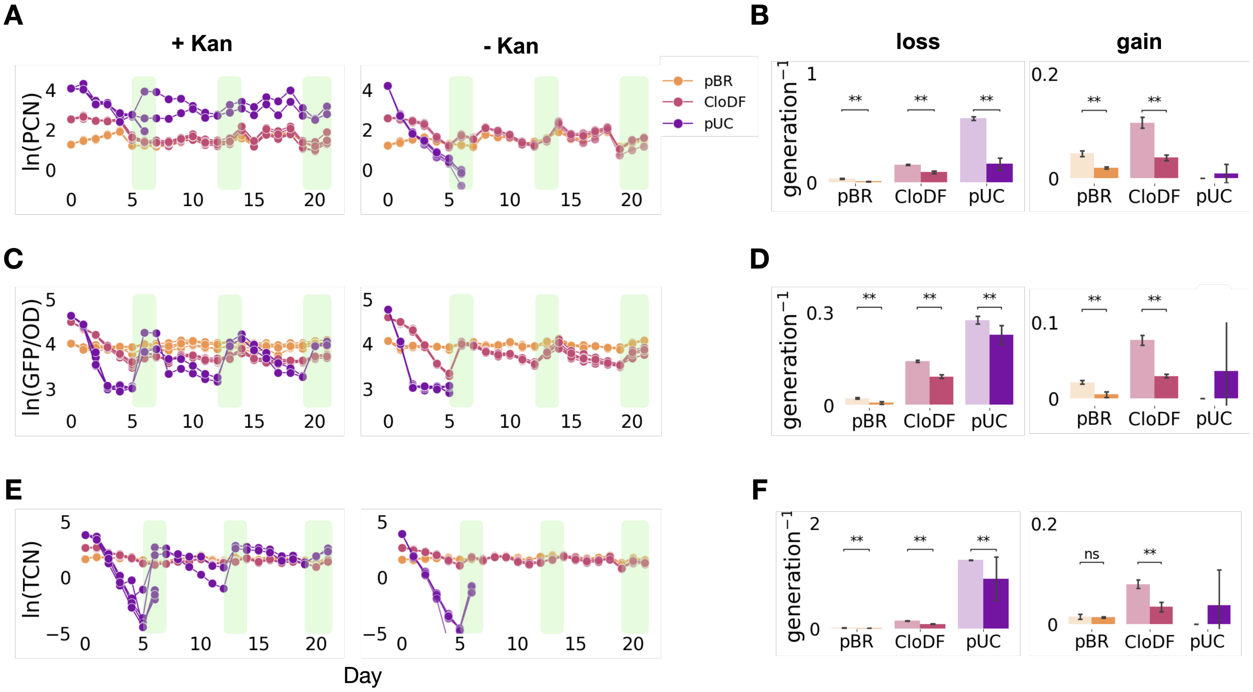


The analyses are the same as in Fig. 4C-H. When there is Kan selection, two out of the five pUC replicates can recover after one course of no transposon selection. When there is no Kan selection, none of the five pUC replicates can recover.

A. qPCR results of the plasmid copy number (PCN) for both conditions, under constant Kan selection and under no Kan selection, of all three plasmids. Left panel shows that constant application of kanamycin maintains the PCN in a narrow range. Right panel shows that PCN varies more drastically without kanamycin selection. The identities of the plasmids are indicated by the three different colors. Legend is on the top right corner.

B. Logarithm fold change of the PCN for both experiments. Left panel is for the loss rate and right panel is for the gain rate. Dark color indicates results with Kan selection. Light color indicates results without Kan selection. Dark bars are lower than light bars, except for the gain rates of pUC, as there is no gain rate data for pUC when there is no Kan selection.

C. OD-normalized GFP results for both experiments of all three plasmids. Data is presented in the same arrangement as in panel A.

D. Logarithm fold change of the OD-normalized GFP for both experiments. Data is presented in the same arrangement as in panel B. Dark color indicates results with Kan selection. Light color indicates results without Kan selection. Dark bars are lower than light bars, except for the gain rates of pUC, as there is no gain rate data for pUC when there is no Kan selection.

E. qPCR results of the transposon copy number (TCN) for both experiments of all three plasmids. Data is presented in the same arrangement as in panel A.

F. Logarithm fold change of the TCN for both experiments. Data is presented in the same arrangement as in panel B. Dark color indicates results with Kan selection. Light color indicates results without Kan selection. Dark bars are lower than light bars, except for the gain rates of pUC, as there is no gain rate data for pUC when there is no Kan selection.

All statistical tests use the Mann-Whitney U test. p-value annotation legend:

ns : 0.05< p <= 1.0; *: 0.01 < p <= 0.05; **: 0.001 < p <= 0.01

**Supplementary Figure 8. qPCR primer characterization data**

**A**

**+ transposase**

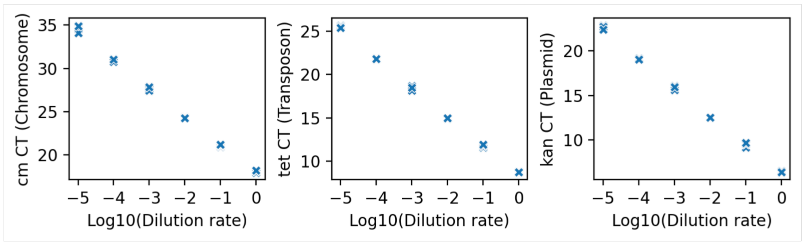


**B**

**- transposase**

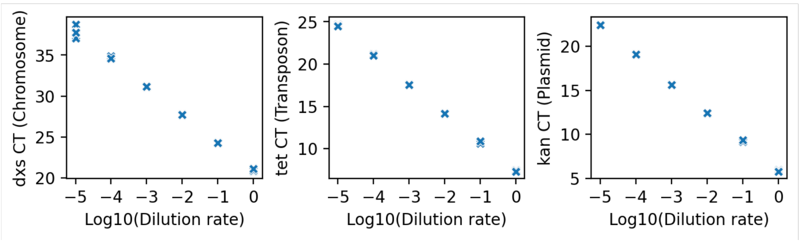


The overnight culture was diluted by mixing 50 µl of culture with 150 µl of molecular-grade water. For cell lysis, 50 µl of the diluted culture was boiled at 98°C for 20 min. Then cell lysate was serially diluted (1/10-fold) and 4 µl of each diluted sample was used in qPCR reactions, with 6 technical replicates performed for each dilution rate. qPCR primers and probes were designed to target the $cm^R$ gene or dxs on E. coli genome (left), the *tetA* gene on transposon (middle) and $kan^R$ gene on the plasmid (right). The $cm^R$ and *dxs* gene is present approximately one copy per genome, enabling for the calculation of the average per-cell pTarget copy number relative to the genome copy number.

To calculate primer efficiencies, Ct values were plotted against log-transformed dilution rates, and a straight line was fitted to the data points.

For cells with transposase, as shown in panel A, the slopes were -3.30 for *cm*, -3.24 for $kan^R$ and -3.35 for *tetA*. The amplification factors were calculated as: $Amplification\ factor = 10^{-\frac{1}{slope}}$ (2.01 for $cm^R$, 2.03 for $kan^R$ and 1.99 for *tetA*). The primer efficiencies were also calculated using: $Primer\ efficiencies = (amplification\ factor - 1) \times 100$ (100.91 % for $cm^R$, 103.47 % for $kan^R$ and 98.97 % for *tetA*).

For cells without transposase, as shown in panel B, the slopes were -3.38 for *dxs*, -3.31 for $kan^R$ and -3.44 for *tetA*. The amplification factors were calculated as: $Amplification\ factor = 10^{-\frac{1}{slope}}$ (1.97 for $cm^R$, 2.01 for $kan^R$ and 1.95 for *tetA*). The primer efficiencies were also calculated using: $Primer\ efficiencies = (amplification\ factor - 1) \times 100$ (97.46 % for $cm^R$, 100.57 % for $kan^R$ and 95.14 % for *tetA*).

**Supplementary Figure 9. Different initial population compositions all lead to positive relationship between PCN and transposon response rate**

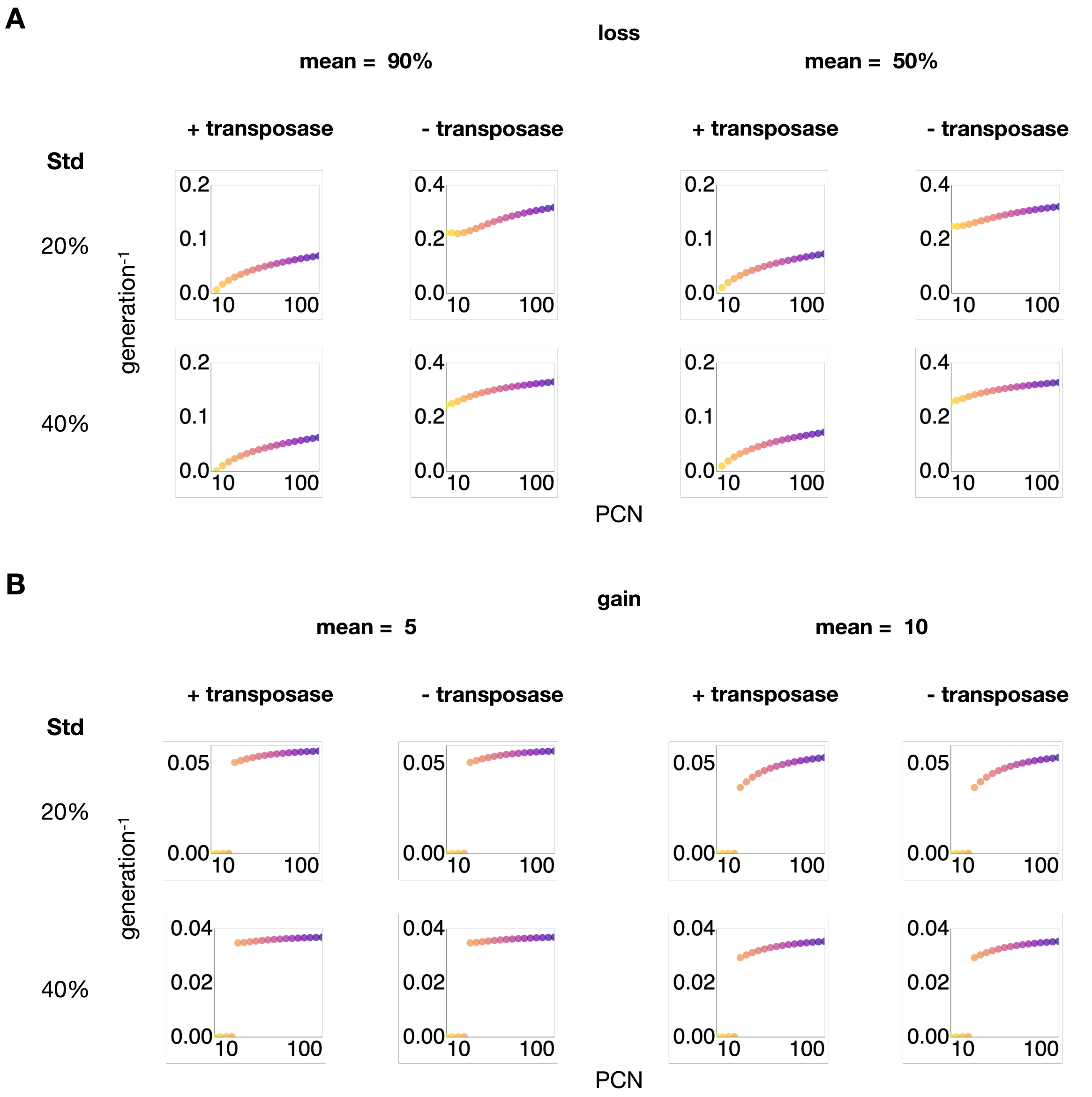


More simulated response rates over PCN, from different initial population composition. Different compositions do not affect the positive relationship between PCN and transposon response rate.

A. Loss rates when there is no selection. The initial composition in this case is a truncated normal distribution with a mean of 90% of maximal TCN (left) or a mean of 50% of maximal TCN (right). The standard deviation is also varied, being either 20% of maximal TCN (top) or 40% of maximal TCN (bottom).

B. Gain rates when there is selection. The initial composition in this case is a truncated normal distribution with a mean of 5 TCN (left) or a mean of 10 TCN (right). The standard deviation is also varied, being either 20% of maximal TCN (top) or 40% of maximal TCN (bottom).

**Supplementary Figure 10. Nonmonotonic relationship between TCN and growth rates lead to nonmonotonically increasing relationship between transposon gain rate and PCN**

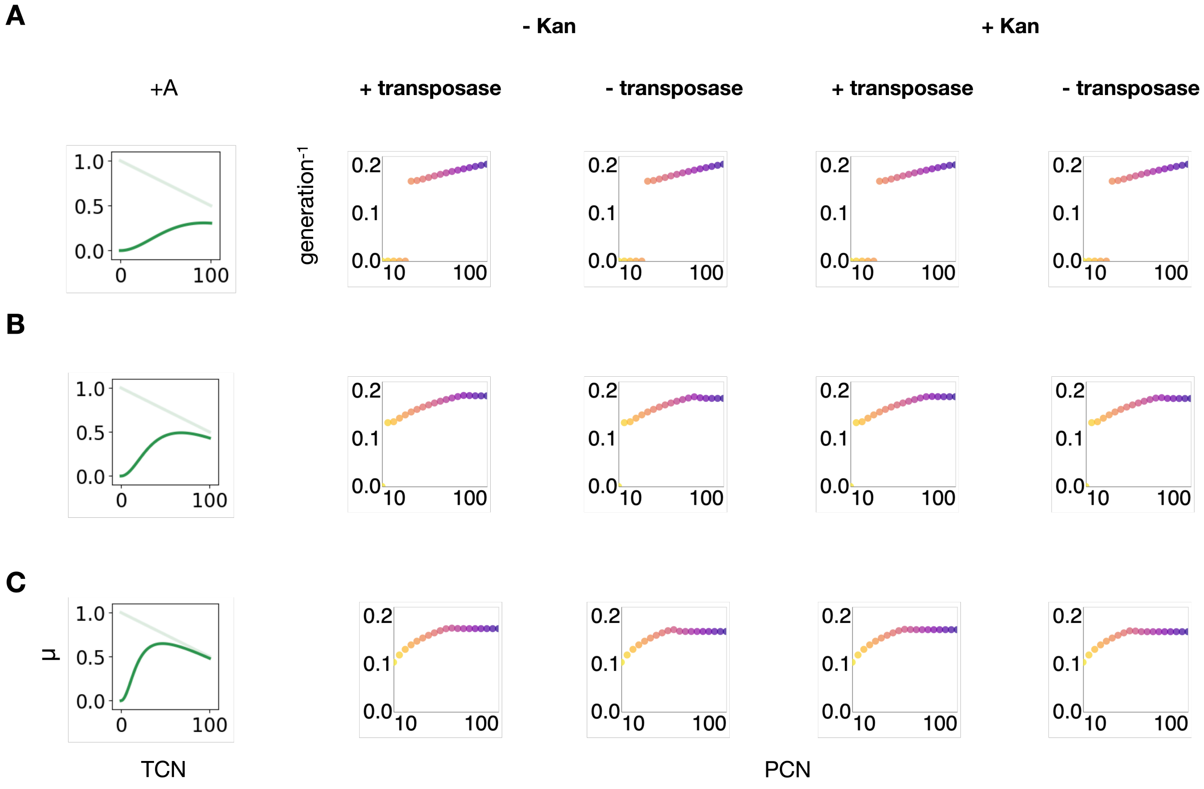


Each row contains simulation results of transposon gain rates under four different conditions, given the growth rate curve on the left side of the row.

A. If growth rate increases monotonically as TCN increases, gain rate increases as PCN increases.

B. If growth rate increases first as TCN increases, then drops as TCN increases further, gain rate increases first, then stops further increasing.

C. Same as panel B but with more drastic nonmonotonic relationship between growth rate and TCN.
